# Generation of natural killer and myeloid cells in a 3D artificial marrow organoid system

**DOI:** 10.1101/2024.01.15.575527

**Authors:** Bérénice Schell, Lin-Pierre Zhao, Camille Kergaravat, Emilie Lereclus, Maria Kalogeraki, Pierre Fenaux, Lionel Ades, Antoine Toubert, Marion Espeli, Karl Balabanian, Emmanuel Clave, Nicolas Dulphy, Valeria Bisio

## Abstract

The human bone marrow (BM) microenvironment involves hematopoietic and non-hematopoietic cell subsets organized in a complex architecture. Tremendous efforts have been made to model it in order to analyse normal or pathological hematopoiesis and its stromal counterpart. Herein, we report an original, fully-human *in vitro* 3D model of the BM microenvironment dedicated to study interactions taking place between mesenchymal stromal cells (MSC) and hematopoietic stem and progenitor cells (HSPC) during the hematopoietic differentiation. This artificial marrow organoid (AMO) model is highly efficient to support NK cell development from the CD34+ HSPC to the terminally differentiated NKG2A-KIR2D+CD57+ NK subset. In addition, myeloid differentiation can also be recapitulated in this model. Moreover, mature NK cell phenotype showed significant differences in the AMO compared to a conventional 2D coculture model for the expression of adhesion molecules and immune checkpoint receptors, thus better reflecting the NK cell behaviour in the BM microenvironment. Lastly, we proved that our model is suitable for evaluating anti-leukemic NK cell function in presence of treatments. Overall, the AMO is a versatile, low cost and simple model able to efficiently recapitulate hematopoiesis and granting better drug response taking into account both immune and non-immune BM microenvironment interactions.

**Graphical Abstract:** 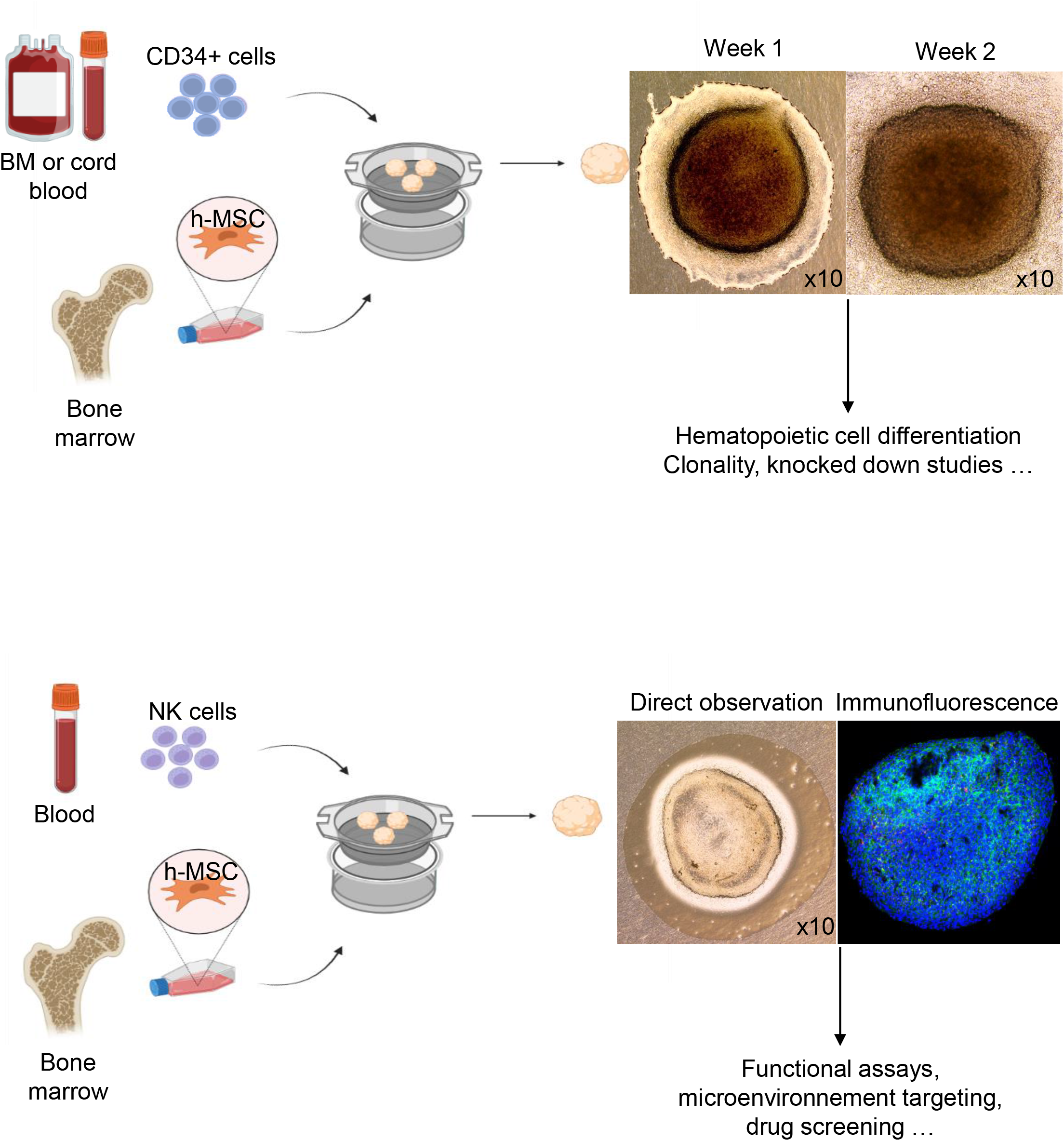

## Introduction

No perfect *in vitro* model has been established so far to study the complex human hematological cell biology. Such a model should be based on the use of primary cells and encompass within the appropriate local bone marrow (BM) microenvironment (BMM) including a functional immune system. Moreover, ideally such model should be easy to use, fast, reproducible and cheap. A wide range of *in vivo* models have been developed but all face some major drawbacks in term of insufficient transferability to human beings (no immunological context), the poor engraftment of hematopoietic stem and progenitor cells (HSPC) into recipient mice and high technical complexity^1,2^.

Therefore, 2D *in vitro* models are still widely used, due to their relative simplicity, cheapness, reproducibility but missing the spatial organization and biomechanical forces of the BMM, crucial for the cellular fate. To overcome this point, there has been a surge of interest for the development of several 3D models^3–5^ that so far do not meet simultaneously the requirements of user-friendliness, reproducibility, low cost and system versatility.

In parallel, the field of Natural Killer (NK) cell-based cancer therapy has grown exponentially and currently constitutes a major area of immunotherapy innovation^6^. NK-cells are a heterogeneous subset of cytotoxic lymphocytes, produced within the BM from the HSPCs, and functionally regulated by a repertoire of activating and inhibitory receptors. Through the production of cytokines (IFN-γ/TNF-α) and cytolytic proteins (perforin/granzymes) and the death receptors, NK-cells are very potent to eliminate cancer cells^7^. Consequently, various strategies to generate NK-cells for adoptive immunotherapy have been developed, mostly relying on a combinations of cytokines with or without feeder cells^7–9^, each with its specific advantages and disadvantages in regard of cell numbers, function and handling efforts. Yet, how the BMM can support, reshape and eventually interfere with the production and function of NK progenitor and mature cells remains largely unknown.

Here, we describe an easy-to-access, reliable and reproducible self-organizing *in vitro* 3D model of human BMM for studying progenitor and mature NK cells in the frame of their physical crosstalk with the human mesenchymal BM stroma.

## Results

### NK and myeloid cell development in the 3D *in vitro* model

The Artificial Marrow Organoid (AMO), is based on the *Seet et al*. model^10^ in which we substituted the murine stromal cell line MS5-hDLL1 with *ex vivo* expanded human primary MSCs (h-MSC). In the model, the MSC conserved mesenchymal-lineage markers (CD73+CD90+CD106+), formed compact clusters with fibrous connections, and retained lineage differentiation capacity (Supplemental Figure 1). Importantly, this self-organizing model does not need any chemical or mineral support by contrast to other systems.

Taking advantage of the simplicity and high reproducibility of this model, we tested whether it could support NK-cell development starting from freshly isolated human cord blood CD34+ HSPCs. H-MSCs (n=150,000), sorted by cell-to-plastic adhesion of BM samples from hip replacement surgery, were co-cultured with a small number of sorted CD34+ HSPCs (n= 6,000/7,500) in presence of a specific cocktail of cytokines (Supplemental Figure 1A). NK-cell commitment was completed within 1 to 4 weeks and up to 2x10^5^ total NK-cells were generated. Flow-cytometry analysis, using an appropriate gating strategy (Figure 1A), based on previously described NK-cell maturation stages^11^ (Supplemental Figure 2A) and pseudotime trajectories (Figure 1B), were used to visualize the progression of HSPCs from immature into various NK maturation stages. In details, already from week 2, the AMO supported efficient NK-cell lineage commitment from CD34+CD38-hematopoietic stem cells (HSCs), as shown by a predominance of CD3-CD56+CD16+/-cells among the CD45RA+ hematopoietic cells (Figure 1A). The AMO model recapitulated the full NK-cell differentiation starting from HSCs (stage 1) to common lymphoid progenitor (CLP, stage 2), Lin-CD56-(stage 3), Lin-CD56+CD16-(stage 4), CD56+CD16+ (stage 5) and CD56+CD16+CD57+ (stage 6) NK-cells (Figure 1A-B). Pseudotime analysis recapitulates this maturation trajectory from CD34+ HSC and CLP with a progressive acquisition of CD56+ and others NK markers such as CD94 and NKp46, acquired simultaneously during stage 3, and NKp80 acquired at stage 4 (Figure 1B and Supplemental Figure 2B). Very few immature T (CD7+CD56-) and mature B cells (CD19+) originate from HSPC under this differentiating protocol. The supervised population gating showed that, by week 2, the majority of cells have engaged into NK-cell differentiation (stage 3: 47.9% and stage 4: 41.3%) while only few HSC (0.3%) and CLP (2.1%) were present (Figure 1C). These immature cells completely disappeared by week 4 where the majority of cells belong to stage 4 (77.2%) with an increase of cells in stage 5 (13.5%). As expected, the NK stage 5 was characterized by the expression of activating (NKp30/DNAM-1) and inhibitory receptors (NKG2A/KIR molecules) crucial for NK-cell functionality (Figure 1D). By contrast to *in vitro* 2D co-culture^7,9^, the AMO is not only able to support NK-cell differentiation but also to reproduce the natural NK-cell heterogeneity (according to the expression of NKG2A and KIR molecules), until the terminally differentiated NKG2A-KIR+CD57+ NK-cell subset (stage 6). Importantly, NK-cells produced after AMO-based differentiation were polyfunctional, as demonstrated by their capacity to degranulate cytolytic granules and produce cytokines (Figure 1E, Supplemental Table 1). In fact, the proportion of NK-cells with monofunction, bifunction and trifunction significantly increased after stimulation by K562 cell line or PMA and ionomycin. Remarkably, the cytotoxic capacity against K562 cells, albeit variable, tended to be higher than that of mature NK-cells sorted directly from peripheral blood (n=3, Supplemental Figure 2B), thus confirming that this model allowed the production of functional and fully differentiated NK-cells with a high efficiency.

**Figure 1.**
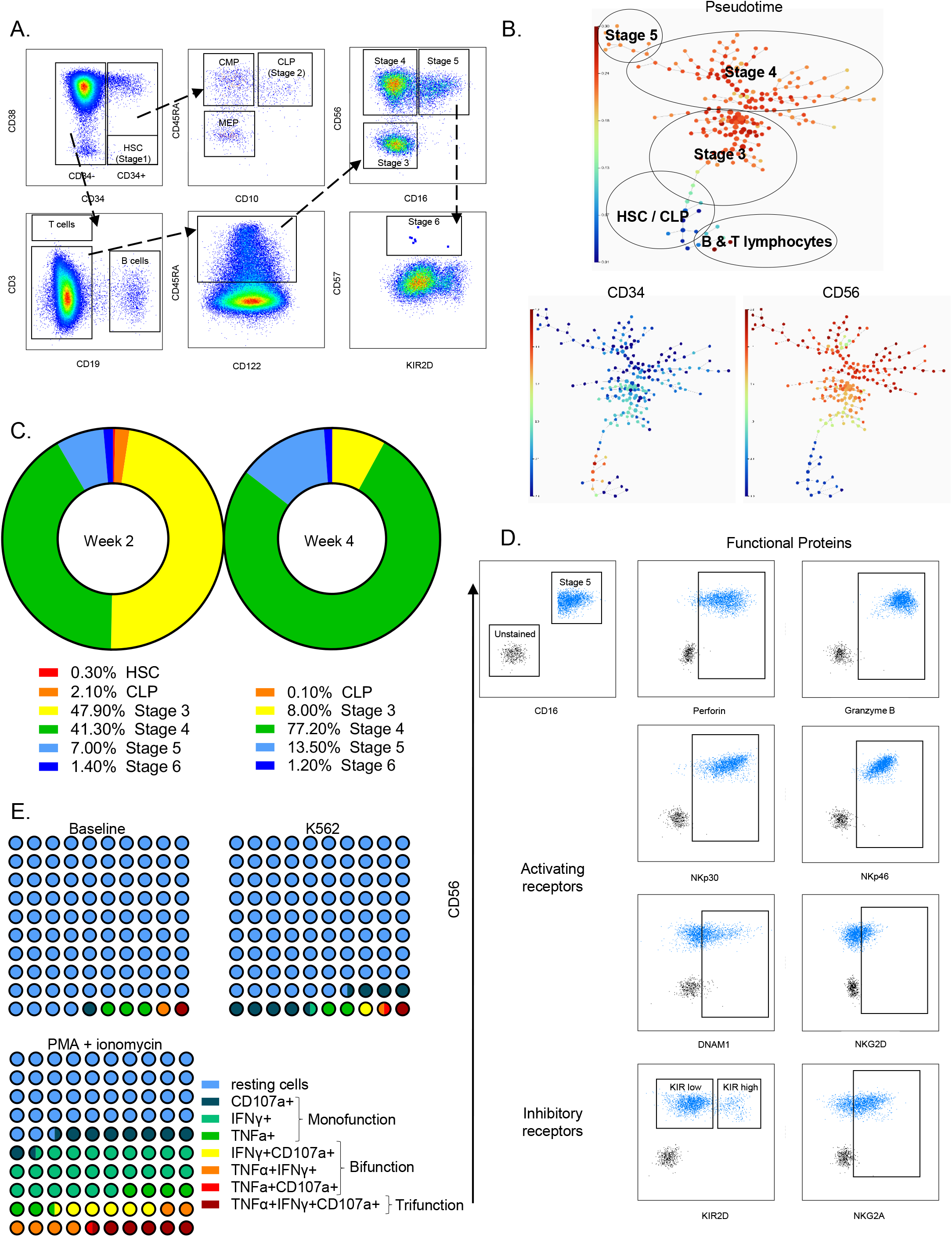
NK cell differentiation from HSPC to mature effector cells. (**A**) Gating strategy allowing to distinguish the different stages of NK cell maturation: HSC (CD34+ CD38-), CLP (CD34+ CD38+ CD10+ CD45RA+), immature NK cells (stage 3: CD34-CD45RA+ CD56-CD16-), and mature NK cells (stage 4: CD34-CD45RA+ CD56bright CD16-, stage 5: CD34-CD45RA+ CD56dim CD16+ and stage 6: CD34-CD45RA+ CD56dim CD16+ CD57+ KIR2D+/-). Plots at two weeks of differentiation. (**B**) Tree representation of SPADE clustering colored by pseudotime calculated using trajectory analysis with Wishbone algorithm of NK cell differentiation taking CD34+CD38-HSC population as starting cell type. The size of nodes is proportional to the number of cells in the given cluster. Calculation and figure were made with the OMIQ software (https://www.omiq.ai/). The differentiation stages have been manually added to the figure in base of the expression of main lineage markers. The same SPADE tree is represented in the lower panel colored by CD34 and CD56 intensity. For other markers, see Supplemental Figure 2B. (**C**) Evolution of NK cell differentiation stages in percentage at two and four weeks of maturation inside the AMO (n=2 at 2 weeks and n=3 at 4 weeks). Gaiting strategy above mention and recalculated to 100%. (**D**) Phenotype of a representative AMO at two weeks of differentiation; density plot of live NK cells gated on stage 5 cell subset as defined in (A). The positivity of NK cell receptors is assessed by comparison to unstained cells gated on forward scatter. (**E**) Evaluation of NK cell’s polyfunctionality after 3 weeks of differentiation inside the AMO (n=4). After organoid dissociation, NK cells were cultured either in presence of the NK-sensitive K562 cell line or with PMA-Ionomycin. NK cell’s degranulation of cytolytic granules (surface expression of CD107a) and IFN-γ and TNF-α intracellular richness were determined by flow cytometry. Proportion of NK-cells with monofunction, bifunction and trifunction significantly increased after stimulation by K562 cell line or PMA and ionomycin (p-value of 0.02 and <0.01 for K562 and PMA-Ionomycin versus control respectively, Fisher exact test).

In parallel, to verify the versatility of the system, we tested the ability of the AMO to recapitulate another cell fate process, the myeloid differentiation. The AMO, setup as described above, was cultured in a different cocktail (of cytokines see methods for details) promoting myeloid differentiation. In this case, the CD34+Lin-HSPCs progressively differentiated loosing early myeloid markers such as CD34, CD117 and HLA-DR, and acquiring CD45 and CD15 overtime (Supplemental Figure 3A). The typical biphasic evolution of CD13 and CD33 with a maximum expression at the promyelocyte stage (here achieved in around 7 days) was in accordance with the myeloid marker evolution already described^12^. The trajectory-based analysis performed at one and two weeks individualized both monocytic and neutrophils branches containing CD64+CD14+ and CD15+CD13+ cells respectively (Supplemental Figure 3B). All this was confirmed by cytological evaluation (Supplemental Figure 3C). Altogether, these data demonstrated the versatility of the AMO to recapitulate innate hematopoiesis, as demonstrated by NK-cells or myeloid subsets generation, by varying only the cytokines cocktails within a primary human MSC environment.

### Mature NK cells immunomodulation in the AMO system

Finally, to verify if the model may reproduce the immunomodulating effect of the BM stroma over immune cells^13^, mature NK-cells isolated from PBMC were co-cultured with h-MSC in the AMO model (Supplemental Figure 1B) and compared to the standard 2D co-culture. Histological analysis showed the infiltration of isolated NK-cells, surrounded by MSC within the 3D structure (Figure 2A). After three days of co-culture of the same MSC and NK-cells in either 2D or AMO systems at the same MSC/NK ratio, the multidimensionality reduction performed on extracted MFIs (mean fluorescence intensity) on gated NK-cells showed dissimilarities in NK-cell phenotype according to the co-culture model (Figure 2B). Contrary to lineage markers that showed a conserved expression on NK-cells co-cultured in 2D and AMO, we found that some adhesion markers, as integrin beta chains 1 and 4 (CD18 and CD49d respectively), were downregulated in the AMO compared to 2D co-culture (Figure 2C). We hypothesized that this downregulation could be due to a higher engagement of these particular integrins as 3D co-cultures favor tighter contacts between cells. We also noticed significant different expression for the immune-checkpoint markers CD73 and Tim-3 and wondered how this could affect NK-cell functionality. Interestingly, NK-cells co-cultured in AMO displayed reduced proliferation and cytotoxic activity compared to the 2D model (Figure 2D), prompting us to conclude that AMO co-cultured MSC displayed higher immunomodulatory properties, as already shown by enhanced pro-survival and anti-inflammatory properties in other spheroid systems^14,15^ and closer to the one described in BM NK-cells^16^. These results suggested that the AMO model could more accurately reproduce the immunoregulations provided by the microenvironment within the BM. Therefore, we explored whether the AMO may be suitable to test drug *in vitro*. First, the AMO cultured in presence of the classical chemotherapy doxorubicin showed a good drug delivery and permeability in the whole structure as shown by fluorescence measure on confocal imaging compared to DMSO alone (Figure 2E). This result prompted us to test treatments with the hypomethylating agents azacytidine (AZA) and decitabine (5-aza-2’-deoxycytidine, DEC), commonly used to treat high-risk MDS (Myelodysplastic Syndromes) and AML (Acute myeloid leukemia) patients, and known to increase the NK cytotoxic functionality^9,17^. Indeed, we noticed the expected increase of NK cytotoxic confirming the drug efficacy within the AMO system (Figure 2F), in particular after DEC treatment, more markedly in AMO system than in 2D. Overall, the AMO system probably better reflects BM microenvironment interactions appearing more suitable for *in vitro* drug testing.

**Figure 2.**
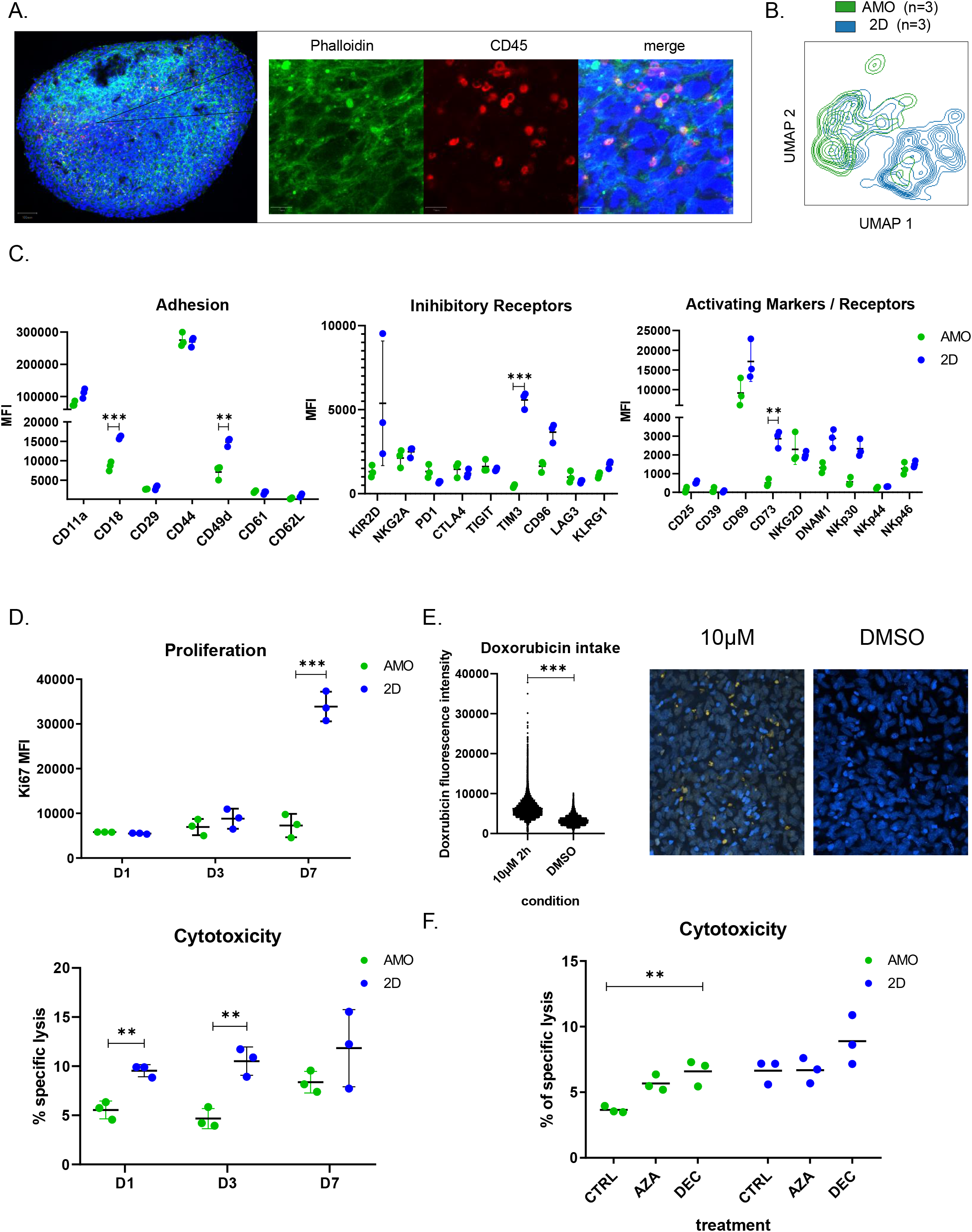
Study of immunomodulating microenvironment and drug targeting. (**A**) AMO observed by confocal imaging after staining with Hoescht (nuclei, blue), Phalloidin-FITC (actin filaments, green) and CD45-AF647 (NK cell membrane, red). Entire organoid in the left panel. NK cell infiltration (in red) in the right panel. X400 magnification. One representative experiment of three is presented. (**B**) UMAP representation of spectral flow-cytometry analysis performed on NK cells from the same donor co-cultured for three days with h-MSC (n=3) in AMO (green) or 2D (blue) cultures. (**C**) Dot plots showing the median of fluorescence (MFI) for main adhesion, inhibitory and activating markers on mature NK cells assessed by spectral flow-cytometry after three days of AMO (green) or 2D (blue) cultures with h-MSC (n=3). Paired Mann-Whitney test, p-value < 0.05. (**D**) Overtime assessment of proliferation (Ki67 MFI, upper panel) and cytotoxicity against K562 cell line (measure of calcein release by fluorescence and expressed in percentage of specific lysis based on spontaneous and maximum release, lower panel). (**E**) Confocal quantification and imaging on the AMO model after treatment with doxorubicin (10μM for two hours) vs control (DMSO). Hoescht in blue and Doxorubicin in yellow, x400 magnification. (**F**) Quantification of the cytotoxic activity measured by calcein-release by K562 cell line in presence of NK cells co-cultured in AMO in presence of AZA or DEC for three days (AMO in green and 2D in blue, n=3).

## Discussion

Herein, we illustrated the simplicity of the AMO model to study cell NK and myeloid ontogenesis within an immunomodulatory microenvironment allowing better evaluations of the immunological response and drug targeting. The advantages of this system over the others are (i) the presence of primary human MSC, which create an *in vivo*–mimicking environment for the hematological cells, (ii) the possibility to study different hematological cells changing only the cocktail of cytokines, (iii) the 3D context granting better prescreening data. Further studies complexifying the AMO with other cell types (as immune CD8+ T, endothelial or leukemic cells) or in autologous setting could further increase the reliability of the model. Meanwhile, the AMO is a straightforward, versatile, reproducible and scalable platform meeting basic and clinical exigencies.

## Methods

### Primary cells

Bone marrow (BM) samples were obtained from patients (ages 18– 80), free of hematological disease and coming to the Orthopedic and Trauma surgery Department of Lariboisiere Hospital (Paris, France) for a total hip replacement surgery. Cord blood (CB) and peripheral blood samples were provided from the blood biobank of Saint-Louis Hospital (Paris, France). All the patients provided written informed consent, in accordance with the Helsinki Declaration. Approval from the Institutional Review Board Paris-Nord was obtained before the use of the clinical materials for research purposes. White cells were purified from the BM or blood samples with a Ficoll gradient (Pancoll Human, PAN-BIOTECH). Human CD34+ cells were magnetically enriched with the EasySep™ Human CD34 Positive Selection Kit II (STEMCELL Technologies) from either normal BM or CB samples. In the same way, mature NK cells were isolated from peripheral blood mononucleated cells with the EasySep™ Human NK Cell Enrichment Kit (STEMCELL Technologies). The human mesenchymal stromal cells (h-MSC) were derived from BM samples and separated from other hematopoietic cells through their plastic adherence after 2-3 weeks of *in vitro* culture. H-MSC were then expanded in αMEM (Gibco) supplemented with 10% fetal bovine serum (FBS, Sigma), 2 mM glutamine, 100 U/mL penicillin and 100 μg/mL streptomycin (Gibco). When the h-MSC reached the 80-90% confluence, the cells were detached with Trypsin (Gibco) and cryopreserved before use. H-MSC were used within 5 passages.

### Artificial Marrow Organoid (AMO) culture

The 3D culture was adapted from *Seet at al*.^10^. In details, h-MSC were harvested by trypsinization and resuspended in serum-free culture medium, so-called AMO medium, composed of RPMI 1640 (Gibco), 4% B27 supplement (ThermoFisher Scientific), 30 μM L-ascorbic acid 2-phosphate sesquimagnesium salt hydrate (Sigma) reconstituted in PBS, 100 U/mL penicillin and 100 μg/mL streptomycin (Gibco), 2 mM glutamine (ThermoFisher Scientific) and a precise cocktail of cytokines depending on the experiments and constituted as follows: myeloid differentiation cytokines’ cocktail: IL-3 7 ng/mL (Miltenyi), SCF 100 ng/mL (Miltenyi), TPO 10 ng/mL (Miltenyi), FLT3 200 ng/mL (Miltenyi); NK differentiation cytokines’ cocktail: IL-7 10 ng/mL (Miltenyi), SCF 10 ng/mL (Miltenyi), TPO 10 ng/mL (Miltenyi), FLT3-L 10 ng/mL (Miltenyi), IL-15 20 ng/mL (Miltenyi), SR-1 2 μM (Miltenyi); NK immunomodulation cocktail: IL-2 100 UI/mL (Miltenyi). Depending on the experiment, different amount of h-MSC cells, from 8×10^4^ to 1.5×10^5^, were combined with 6×10^3^–7.5×10^3^ purified CD34+ cells or 4×10^4^ NK cells per AMO in 1.5 ml Eppendorf tubes and centrifuged at 300 g for 5 min. Supernatants were carefully removed and the cell pellet was harvested. For each AMO, a 0.4 μm Millicell transwell insert (EMD Millipore) was placed in a 6-well plate containing 1 mL of AMO medium per well supplemented with the appropriate cytokines. Medium was changed completely every 3 days. The AMO were kept in culture for different time points depending on the experiments: up to two weeks for the myeloid maturation, three weeks for the NK ontogenesis and 1 to 7 days maximum for the NK immunomodulatory experiments. At the indicated time points, AMOs were harvested and dissociated in collagenase (Thermofisher) at a final concentration of 200 U/mL in 1X PBS for 30 minutes at 37°C. If the analysis of the MSC is not required, a passage through a 50 μm nylon strainer is recommended.

### 2D cell culture

MSCs were seeded overnight on 24 or 6 well plates in normal α-MEM supplemented with 10% fetal bovine serum (FBS, Sigma), 2 mM glutamine, 100 U/mL penicillin and 100 μg/mL streptomycin (Gibco). The day after, the media was removed and the NK cells were added to the culture conditions (at the same ratio NK:MSC used in the AMO culture). The co-cultures were maintained in the same AMO medium and analyzed in parallel with the 3D systems.

### H-MSC differentiation

To evaluate differentiation and clonogenic capacities of h-MSC after 3D culture, AMOs were dissociated and cells were counted. CFU-F (colony forming units-fibroblast), adipogenic and osteogenic differentiations were performed as described elsewhere^18^. Briefly, for CFU-F, 200 h-MSCs were distributed in 25 cm^2^ flask in 7mL of complete α-MEM FBS medium for three weeks with a medium change once a week. 5.000 h-MSCs were distributed in 24-well plates for osteogenic and adipogenic differentiation for 21 days. After 24h of adhesion, the osteogenic or adipogenic media were added onto the induced wells, whereas complete α-MEM FBS medium or adipogenic maintenance medium was added onto the control wells. At the end of the experiments, medium was removed, cells were fixed and stained following fabricant’s recommendations with 2% crystal violet (SIGMA), 2% red alizarin (SIGMA) or oil red (Lipid oil red staining kit, SIGMA) to assess CFU-F, osteogenic and adipogenic differentiation respectively (Supplemental Figure 1 D-F).

### Functional assays on NK cells

The cytotoxicity of NK cells was assessed by a calcein release assay. The NK-sensitive K562 cell line was labeled with 1 μg/mL calcein (ThermoFisher Scientific) in PBS for 1h at 37°C. NK cells were dissociated from the AMO, enumerated and co-cultured with calcein-labeled K562 cells in 96-well U-bottom plates in a medium supplemented with Probenecid at 1:1 effector:target ratio. The killing was quantified after 4 h of incubation at 37 °C by measuring calcein-release into the supernatant. The specific killing was calculated as follows: (Measured fluorescence of K562 + NK cell well—Spontaneous Fluorescence)/(Maximum fluorescence—Spontaneous fluorescence)*100. For the IFN-γ and TNF-α quantification, enriched NK cells were cultured with PMA-Ionomycin (SIGMA, 50 and 500 ng/mL, respectively) or K562 cells (1:1 effector:target ratio) for 5 h at 37 °C. Brefeldin A (Sigma) was added at a final concentration of 10 μg/ml after 1 h of incubation. The percentage of CD107a, IFN-γ and TNF-α positive cells was estimated by flow cytometry in the CD3−CD56+ NK cells. Spontaneous release was detected in the absence of target cells.

### Flow-cytometry analysis

After AMO dissociation, cells were washed in PBS and stained according to different panels presented in the Supplemental Table 2. Supervised analyses were performed on data extracted from FlowJo v10.7 software and analyzed with Graph Pad Prism v8.0 software. Uniform manifold approximation and projection (UMAP) and trajectory analyses were performed with OMIC software from Dotmatics (www.omiq.ai, www.dotmatics.com).

### Staining and microscopy

The AMO were fixed, without prior dissociation, with 4% paraformaldehyde (PFA, Sigma) for 15 minutes and permabilized with 0.3% Triton-X-100 (Sigma) in PBS for 15 minutes. Blocking solution (0.5% BSA in PBS) was then used for at least 1 hour. After the blocking step, the AlexaFluor647-labeled CD45-specific monoclonal antibody (Biolegend), FITC-labeled Phalloidin (Sigma) and Hoechst 33342 (ThermoFisher Scientific) were used in a solution with saponin 0.025% in PBS, and incubated with the AMO overnight at 4°C. Images were acquired by laser scanning microscopy on a LSM 800 AiryScan system mounted on an Axio Observer stand (Zeiss, Oberkochen, Germany) and with a Plan Apochromat 63X N.A. 1.4 oil-immersion objective using the Zen Blue edition software (Zeiss, v2.3). The pixel size was set at 0.71 μm and a z-step of 0.5 μm was used. The quantification of doxorubicin fluorescence was performed using QuPath software (v0.3.2). May Grunwald Giemsa (MGG) staining was performed on harvested and cytospined cells after AMO co-culture with myeloid differentiation induction and dissociation. Different stages of the myeloid differentiation were morphologically identified by optical microscopy at x100 magnification and counted over a total of 200 cells.

### *In vitro* cell treatment and functional assay

AMO were treated in AMO cell culture medium with 0.5 μM of AZA, DAC (Sigma). DMSO condition was used as control. The treatments were maintained for 3 days with a half change of medium after 48 h (IL-2 + drugs or DMSO). Then, cells were collected and NK cells were analyzed for their cytotoxic capacity.

## Acknowledgments

This study was supported by a grant from the French Ministry of Health and the French National Cancer Institute (#PRT-K2017-109), the Cancéropôle Ile-de-France, the Association Laurette Fugain (#ALF 2016-07), the Association Force Hémato (Call for projects 2017), the Ligue contre le Cancer (Ile-de-France committee), and the Fondation de France (Call for Basic and Translational Research in Cancer proposals 2020) and the IHU THEMA (Call 2022). The authors thank the Technological Core Facility of the Saint-Louis Research Institute, UMS “Saint-Louis”, US53/UAR2030. The facility is supported by grants from the University de Paris, the Conseil Regional d’Ile-de-France (Canceropôle), the National Cancer Research Institute (InCa), the Ministère de la Recherche, the Association Saint-Louis and the Association Jean Bernard. INSERM UMR 1160 is a member of OPALE Carnot Institute, The Organization for Partnerships in Leukemia, Institut de Recherche Saint-Louis, Hôpital Saint-Louis, Paris, France (www.opale.org).

## Authorship contributions

BS, LPZ, ND and VB designed, analyzed and interpreted all experiments, and wrote the manuscript. BS, LPZ, CK and EL performed experiments. MK analyzed microscopy experiments. PF, LA, AT, ME and KB contributed to the scientific orientation of the study and critically reviewed the manuscript. CK, AT and EC contributed to the design of the experimental model. ND and VB supervised the study.

## Conflict-of-interest disclosure

The authors declare no competing financial interests.

## Notes

### Competing Interest Statement

The authors have declared no competing interest.

