## Supplemental Information for "Generation of natural killer and myeloid cells in a 3D artificial marrow organoid system"

### Supplementary information

#### Supplemental Tables.

**Supplemental Table 1. Polyfunctional Assay after NK differentiation from cord blood CD34+ cells. Expressed as percentage of positive cells among NK cells (n=4)**

| | Resting cells | CD107a+ | IFN-γ+ | TNF-α+ | IFN-γ+ | TNF-α+ IFN-γ+ | TNF-α+ CD107a+ | TNF-α+ IFN-γ+ CD107a+ |
|---|---|---|---|---|---|---|---|---|
| **Baseline** | 94.1 (91.8-95.5) | 1.1 (0.6-1.7) | 0.5 (0.1-1.1) | 2.3 (1.3-2.8) | 0.2 (0.1-0.6) | 0.9 (0.3-1.7) | 0.2 (0.1-0.4) | 0.7 (0.1-1.1) |
| **K562** | 83.4 (79.6-86.4) | 9.2 (6.2-13.2) | 1.9 (0.4-4.9) | 1.4 (0.5-2.1) | 1.4 (0.4-2.1) | 0.7 (0.3-1.1) | 0.7 (0.5-1.4) | 1.3 (0.5-3.0) |
| **PMA + ionomycin** | 37.3 (29.2-42.3) | 11.0 (9.3-14.4) | 26.3 (24.2-29.) | 5.4 (4.1-6.8) | 9.7 (5.3-17.2) | 4.5 (2.8-5.9) | 0.6 (0.5-0.9) | 5.2 (2.9-6.8) |

**Supplemental Table 2. List of antibodies used for flow cytometry**

| **Myeloid differentiation panel** | | | | |
|---|---|---|---|---|
| Anti-human Ab | clone | Isotype | Supplier | Catalog# |
| CD45 | HI30 | Mouse IgG1 | Thermofisher | MHCD4530 |
| CD34 | 581 | Mouse IgG1 κ | Biolegend | 343508 |
| CD38 | HIT2 | Mouse IgG1 κ | BD | 555459 |
| CD33 | WM53 | Mouse IgG1 κ | Biolegend | 303415 |
| CD15 | W6D3 | Mouse IgG1 κ | BD | 747426 |
| CD11b | ICRF44 | Mouse IgG1 κ | BD | 741601 |
| CD16 | 3G8 | Mouse IgG1 κ | BD | 560715 |
| **NK differentiation panel** | | | | |
| CD45RA | HI100 | Mouse IgG2b κ | BD | 740298 |
| CD16 | 3G8 | Mouse IgG1 κ | BD | 560715 |
| CD19 | SJ25C1 | Mouse IgG1 κ | BD | 612916 |
| CD3 | UCHT1 | Mouse IgG1 κ | BD | 612896 |
| CD161 | DX12 | Mouse IgG1 κ | BD | 562615 |
| CD34 | 563 | Mouse IgG1 κ | BD | 746415 |
| CD7 | M-T701 | Mouse IgG1 κ | BD | 563650 |
| CD33 | WM53 | Mouse IgG1 κ | Biolegend | 303417 |
| CD14 | M5E2 | Mouse IgG2a κ | Biolegend | 301831 |
| DNAM-1 | 11A8 | Mouse IgG1 κ | BD | 752666 |
| CXCR4 | 12G5 | Mouse IgG2a κ | BD | 741643 |
| NKG2A | 131411 | Mouse IgG2a κ | BD | 749682 |
| NKp46 | M5E2 | Mouse IgG2a κ | Biolegend | 301832 |
| IL15Ra | JM7A4 | Mouse IgG2b κ | BD | 747701 |
| NKG2D | 1D11 | Mouse IgG1 κ | BD | 747025 |

| | | | | |
|---|---|---|---|---|
| CD57 | NK-1 | Mouse IgM κ | BD | 563896 |
| CD5 | UCHT2 | Mouse IgG1 κ | Biolegend | 300617 |
| CD122 | TU27 | Mouse IgG1 κ | Biolegend | 339011 |
| NKp80 | REA845 | Mouse IgG1 | Miltenyi | 130-112-591 |
| NKp30 | P30-15 | Mouse IgG1 κ | Biolegend | 325231 |
| CD127 | A019D5 | Mouse IgG1 κ | Biolegend | 351363 |
| CD10 | HI10a | Mouse IgG1 κ | Biolegend | 312206 |
| CD56 | QA17A16 | Mouse IgG1 κ | Biolegend | 392427 |
| CD94 | DX22 | Mouse IgG1 κ | Biolegend | 305516 |
| KIR2D | NKVFS1 | Mouse IgG1 κ | Miltenyi | 130-123-710 |
| CD117 | 104D2 | Mouse IgG1 κ | Biolegend | 313249 |
| CD11b | ICRF44 | Mouse IgG1 κ | Biolegend | 301355 |
| CD27 | M-T271 | Mouse IgG1 κ | Biolegend | 356428 |
| CD38 | HB-7 | Mouse IgG1 κ | Biolegend | 356644 |
| CD107a | REA792 | Mouse IgG1 | Miltenyi | 130-111-702 |
| **NK immunomodulation panel: adhesion** | | | | |
| CD7 | CD7-6B7 | Mouse IgG2a κ | Biolegend | 982702 |
| CD11A | HI111 | Mouse IgG1 κ | BD | 565771 |
| CD16 | 3G8 | Mouse IgG1 κ | BD | 563785 |
| CD18 | L130 | Mouse IgG1 κ | BD | 744553 |
| CD27 | M-T271 | Mouse IgG1 κ | BD | 560609 |
| CD28 | CD28.2 | Mouse IgG1 κ | BD | 741168 |
| CD29 | MAR4 | Mouse IgG1 κ | BD | 564565 |
| CD44 | IM7 | Rat IgG2b κ | BD | 561860 |
| CD49d | 9F10 | Mouse IgG1 κ | BD | 750760 |
| CD56 | B159 | Mouse IgG1 κ | BD | 742022 |
| CD57 | HNK-1 | Mouse IgM κ | Biolegend | 359608 |
| CD73 | AD2 | Mouse IgG1 κ | BD | 748585 |
| CD61 | 2C9.G2 | Hamster IgG1 κ | BD | 566227 |
| CD62L | DREG-56 | Mouse IgG1 κ | BD | 562301 |
| CD95 | SA367H8 | Mouse IgG1 κ | Biolegend | 152612 |
| IL6R | UV4 | Mouse IgG1 κ | Biolegend | 352812 |
| CXCR3 | G025H7 | Mouse IgG1 κ | Biolegend | 353716 |
| CXCR4 | 12G5 | Mouse IgG2a κ | Biolegend | 306508 |
| CCR2 | 48607 | Mouse IgG2b | BD | 558406 |
| CXCR7 | 8F11-M16 | Mouse IgG2b κ | Biolegend | 331104 |
| TRAIL-R1 | S35-934 | Mouse IgG1 | BD | 752308 |
| TRAIL-R2 | B-K29 | Mouse IgG1 | BD | 746679 |
| TRAIL-R3 | B-D44 | Mouse IgG1 | BD | 748959 |
| **NK immunomodulation panel: activation** | | | | |
| CD7 | CD7-6B7 | Mouse IgG2a κ | Biolegend | 982702 |
| CD16 | 3G8 | Mouse IgG1 κ | BD | 563785 |
| CD25 | M-A251 | Mouse IgG1 κ | BD | 751331 |
| CD39 | A1 | Mouse IgG1 κ | BD | 567675 |
| CD56 | B159 | Mouse IgG1 κ | BD | 742022 |
| CD57 | HNK-1 | Mouse IgM κ | Biolegend | 359608 |
| CD69 | FN50 | Mouse IgG1 κ | BD | 612817 |
| CD73 | AD2 | Mouse IgG1 κ | BD | 748585 |
| CD96 | NK92.39 | Mouse IgG1 κ | Biolegend | 338415 |
| CTLA-4 | BNI3 | Mouse IgG2a κ | BD | 566917 |
| LAG-3 | 11C3C65 | Mouse IgG1 κ | Biolegend | 369304 |

| | | | | |
|---|---|---|---|---|
| PD1 | EH12.2H7 | Mouse IgG1 κ | Biolegend | 329950 |
| NKG2D | 1D11 | Mouse IgG1 κ | BD | 749932 |
| NKp44 | 44.189 | Mouse IgG2b κ | Thermofisher | 46-3369-42 |
| DNAM-1 | DX11 | Mouse IgG1 κ | BD | 564796 |
| NKG2A | 131411 | Mouse IgG2a κ | BD | 747917 |
| NKp46 | 9E2 | Mouse IgG1 κ | Biolegend | 331919 |
| KIR2D | NKVFS1 | Mouse IgG1 κ | Miltenyi | 130-123-710 |
| TIM-3 | F38-2E2 | Mouse IgG1 κ | Biolegend | 345034 |
| KLRG1 | REA261 | Mouse IgG1 | Miltenyi | 130-120-423 |
| NKp30 | p30-15 | Mouse IgG1 κ | BD | 563385 |
| TIGIT | A15153G | Mouse IgG2a κ | Miltenyi | 372711 |
| **NK immunomodulation panel: Functional assay** | | | | |
| CD56 | B159 | Mouse IgG1 κ | BD | 557747 |
| CD16 | 3G8 | Mouse IgG1 κ | BD | 563785 |
| CD3 | UCHT1 | Mouse IgG1 κ | BD | 562280 |
| KIR2D | NKVFS1 | Mouse IgG1 κ | Miltenyi | 130-123-710 |
| CD107a | H4A3 | Mouse IgG1 κ | BD | 561343 |
| NKG2A | 131411 | Mouse IgG2a κ | BD | 747917 |
| **Intracellular staining** | | | | |
| Perforin | dG9 | Mouse IgG2b | Biolegend | 308129 |
| Granzyme | GB11 | Mouse IgG1 κ | Biolegend | 515407 |
| Ki67 | Ki-67 | Mouse IgG1 κ | Biolegend | 350510 |
| INF-γ | B27 | Mouse IgG1 κ | BD | 557718 |
| TNF-α | MAb11 | Mouse IgG1 κ | BD | 559321 |
| **Viability dye** | | | | |
| eBioscience™ Fixable Viability Dye eFluor™ 506 | | | ThermoFisher | 65-0866-18 |
| ViaDye Red | | | Cytek | SKUR7-60008 |
| Zombie UV | | | Biolegend | 423107 |
| **Microscopy** | | | | |
| CD45 | HI30 | Mouse IgG1 κ | Biolegend | 304020 |
| Phalloidin | | | SIGMA | P5282-.1MG |
| Hoechst 33342 | | | ThermoFisher | H3570 |

#### Supplemental Figures

**Supplemental Figure 1.** Protocol for the Artificial Marrow Organoid (AMO) formation and observation. AMO can be formed by concentrating MSCs and other cell types and depositing a drop of cell concentrate in a membrane placed over cell-culture medium. (**A**) CD34+ hematopoietic stem and progenitor cells isolated from BM or cord blood were differentiated into NK cells in AMO using an appropriate cocktail of cytokines. Direct optical microscopic observations (x100 magnification) of AMO with CD34+ HSPCs after one and two weeks of culture are shown. (**B**) Mature NK cells were isolated from peripheral blood and cultured in AMO for three days. Direct optical microscopic observations (x100 magnification) of AMO with NK cell at two and ten days. MSC characterization after AMO co-culture: (**C**) Histograms showing expression of three mesenchymal lineage markers CD73, CD90 and CD106 assessed by flow cytometry compared to unstained cells gated on forward scatter; (**D**) Colony Forming Unit assay of MSCs after cristal violet staining at x100 magnification; (**E**) Osteogenic differentiation (top) and control (bottom) of MSCs stained with red alizarin at x100 magnification; (**F**) Adipogenic differentiation (top) and control (bottom) of MSCs observed after oil red staining at x400 magnification. One representative experiment of at list three is presented.

**Supplemental Figure 2.** NK Cell Differentiation from CD34+ HSPCs in AMO. (**A**) Schematic representation of NK cell maturation stages used for manual gating. Image based on *Di Vito et al Front Immunol. 2019* (**B**) Tree representation of SPADE clustering colored by different lineage markers. The size of nodes is proportional to the number of cells in the given cluster. (**C**) Quantification of the cytotoxic activity measured by calcein-release (results expressed as percentage of specific lysis) after three weeks of NK cell differentiation inside the AMO system compared to mature NK sorted from PB.

**Supplemental Figure 3.** Myeloid Cell Differentiation from CD34+ HSPCs in AMO. (**A**) Evolution of the main myeloid differentiation markers, quantified overtime by spectral flow-cytometry, after gating alive cells and eliminating doublets. Data were expressed as Median of Fluorescence in log10 scale. The smoothing was performed with Graph Pad Prism v8.0 software (n=2). (**B**) Tree representation of SPADE clustering colored by pseudotime calculated using trajectory analysis with Wishbone algorithm of myeloid cell differentiation taking CD34+CD38- HSC population as starting cell type. The size of nodes is proportional to the number of cells in the given cluster. Calculation and figure were made with the OMIQ software (https://www.omiq.ai/). The differentiation stages have been manually added to the figure in base of the expression of main lineage markers. The same SPADE tree is represented in the right panel colored by CD13, CD15, CD64 and CD14 intensity. (**C**) Quantitative evaluation of one representative myeloid cell differentiation at two time points (4 and 15 days) observed after MGG staining of AMO smears at x100 magnification. One representative experiment out of two is presented.

**Supplemental Figure 1 – Protocol for the Artificial Marrow Organoid (AMO) formation and observation**

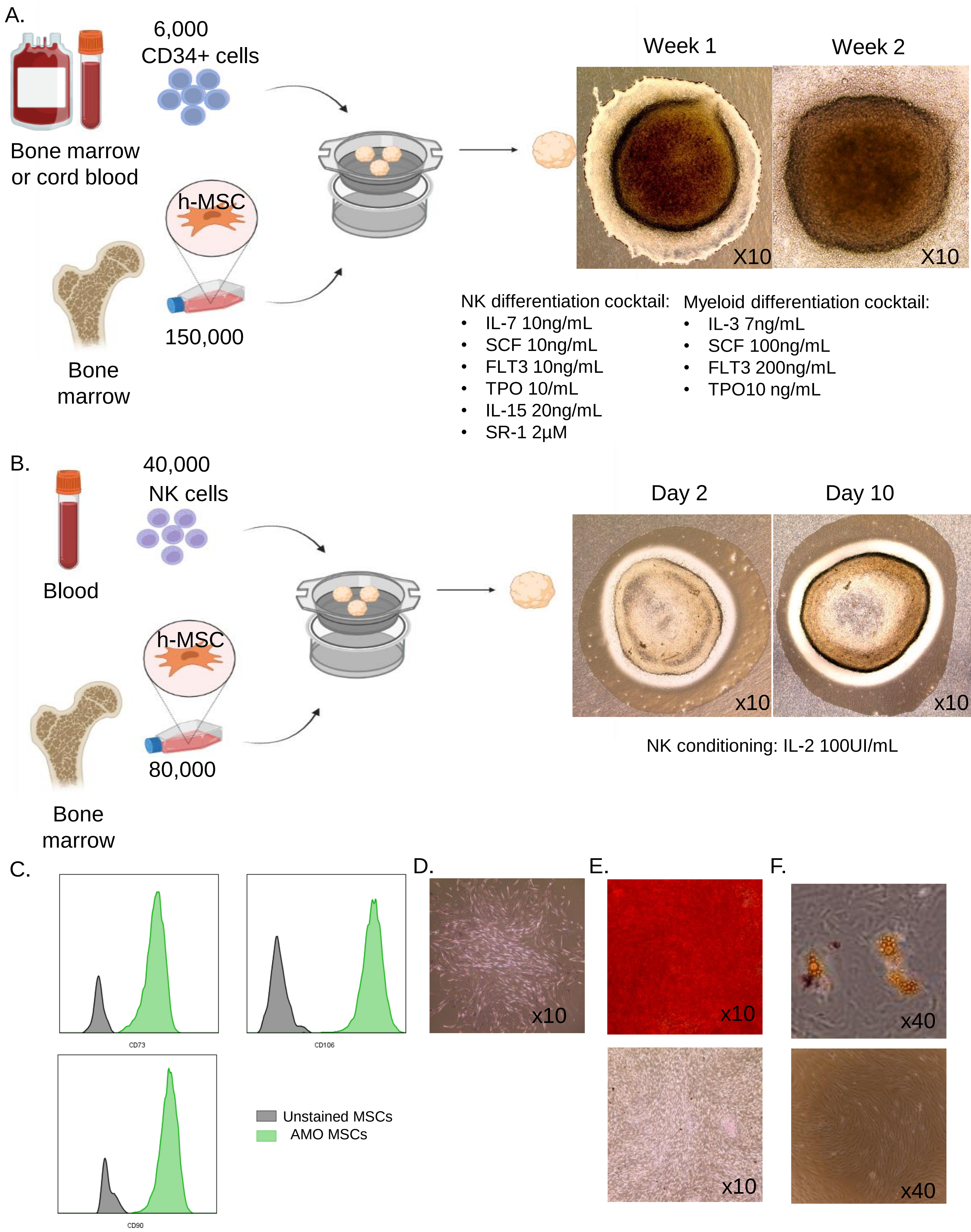

### Supplemental Figure 2 – NK Cell Differentiation from CD34+ HSPCs in AMO

A.

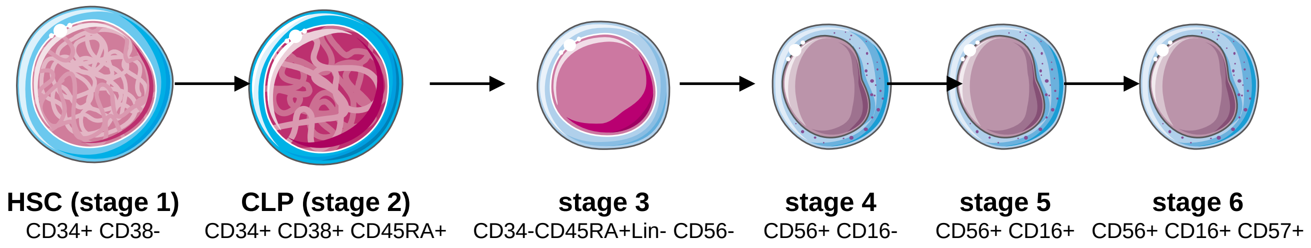


B.

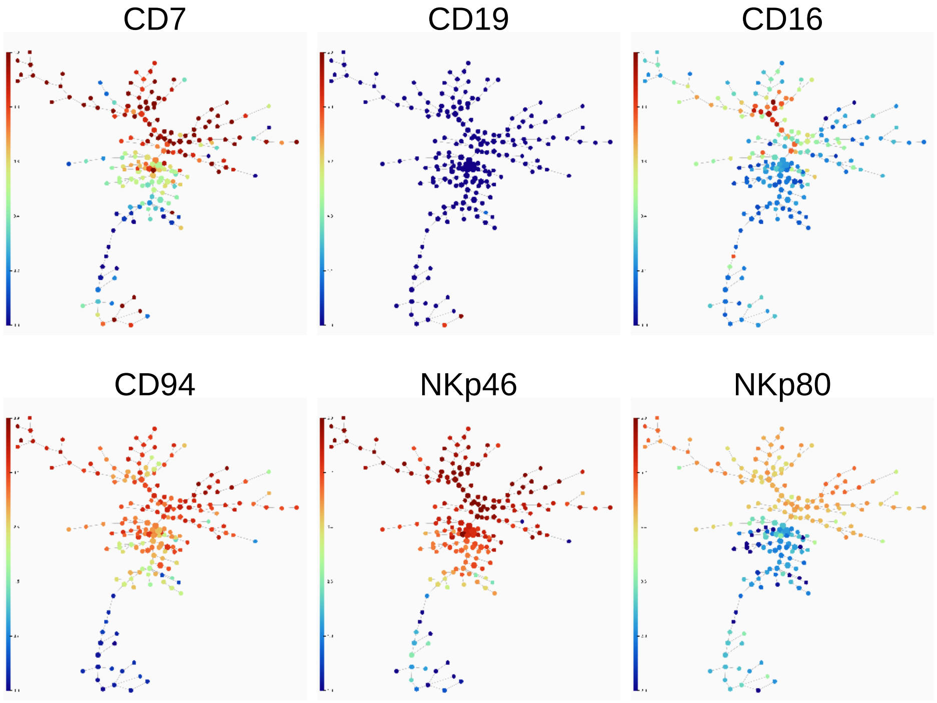


C.

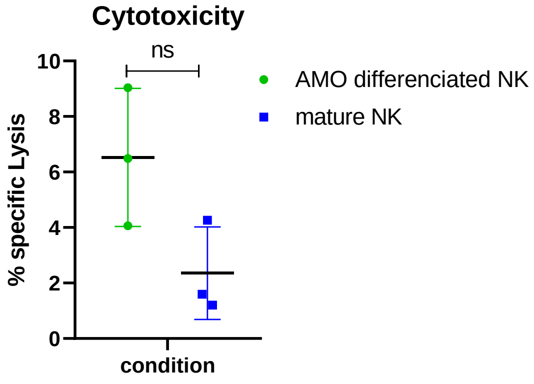

### Supplemental Figure 3 – Myeloid Cell Differentiation from CD34+ HSPCs in AMO

A.

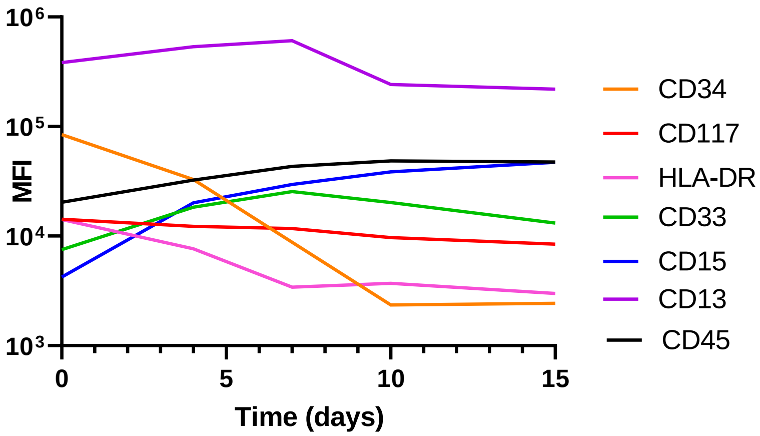


B.

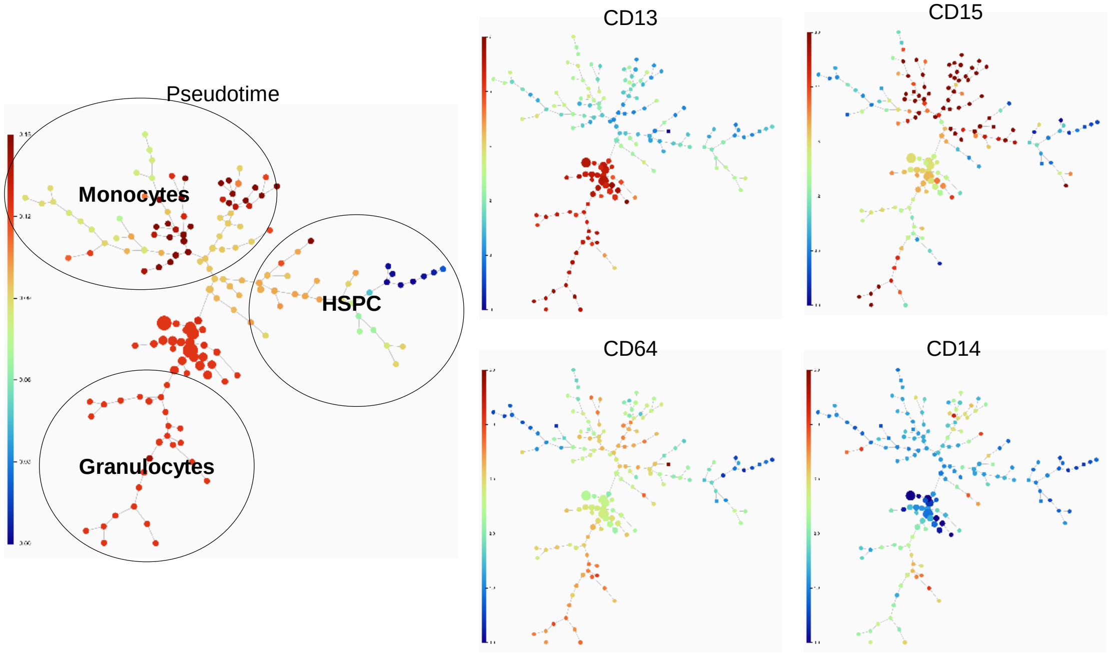


C.

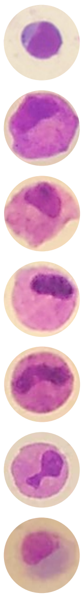

| | D4 | D15 |
|---|---|---|
| Myeloblasts | 32% | 6% |
| Promyelocytes | 14% | 2% |
| Myelocytes | 8% | 12% |
| Metamyelocytes | 6% | 16% |
| Band cells | 12% | 16% |
| Neutrophiles | 12% | 28% |
| Monocytes | 16% | 20% |
